# Transcranial stimulation of alpha oscillations modulates brain state dynamics in sustained attention

**DOI:** 10.1101/2023.05.27.542583

**Authors:** Joshua A. Brown, Kevin J. Clancy, Chaowen Chen, Yimeng Zeng, Shaozheng Qin, Mingzhou Ding, Wen Li

**Affiliations:** Department of Psychology, Florida State University, Tallahassee, FL; Tallahassee Memorial Healthcare, Tallahassee, FL; State Key Laboratory of Cognitive Neuroscience and Learning, Beijing Normal University, Beijing, China; J Crayton Pruitt Family Department of Biomedical Engineering, University of Florida, Gainesville, FL

**Keywords:** complex systems, dynamics, hidden states, non-invasive brain stimulation (NIBS), simultaneous fMRI-tACS

## Abstract

The brain operates an advanced complex system to support mental activities. Cognition is thought to emerge from dynamic states of the complex brain system, which are organized spatially through large-scale neural networks and temporally via neural synchrony. However, specific mechanisms underlying these processes remain obscure. Applying high-definition alpha-frequency transcranial alternating-current stimulation (HD α-tACS) in a continuous performance task (CPT) during functional resonance imaging (fMRI), we causally elucidate these major organizational architectures in a key cognitive operation— sustained attention. We demonstrated that α-tACS enhanced both electroencephalogram (EEG) alpha power and sustained attention, in a correlated fashion. Akin to temporal fluctuations inherent in sustained attention, our hidden Markov modeling (HMM) of fMRI timeseries uncovered several recurrent, dynamic brain states, which were organized through a few major neural networks and regulated by the alpha oscillation. Specifically, during sustain attention, α-tACS regulated the temporal dynamics of the brain states by suppressing a Task-Negative state (characterized by activation of the default mode network/DMN) and Distraction state (with activation of the ventral attention and visual networks). These findings thus linked dynamic states of major neural networks and alpha oscillations, providing important insights into systems-level mechanisms of attention. They also highlight the efficacy of non-invasive oscillatory neuromodulation in probing the functioning of the complex brain system and encourage future clinical applications to improve neural systems health and cognitive performance.

## Introduction

The human brain is an advanced complex system, and two mechanisms—large-scale neural networks and long-range oscillatory neural synchrony —are thought to serve as the primary (spatial and temporal, respectively) organizational architectures of this system^1–4^. Important insights into this organization in the human brain have emerged from functional magnetic resonance imaging (fMRI) and electro/magnetoencephalography (EEG/MEG) research through the identification of reliable intrinsic connectivity networks (such as the default mode network/DMN) and robust canonical oscillations (such as the alpha oscillation). Importantly, growing fMRI and EEG/MEG evidence converges to support the inherent synergy between the two organizational architectures such that the integration of large-scale neural networks both mediates^5^ and is mediated by synchronized oscillations over multiple frequency bands^1,3,4^.

Characteristic of a complex system, the brain is highly dynamical. The past few years have witnessed a major advance in characterizing the spatiotemporal dynamics of the brain in general and the large-scale networks and long-range synchrony specifically^2,6^. Beyond conventional analyses that assume stationarity and provide static (time-averaged) depictions of neural networks and oscillations, this rapidly developing research has identified transient and non-stationary recurring patterns of organized activity in the brain (known as “brain states”), both at rest and during task performance, and demonstrated their relevance to cognition and neuropsychiatric disorders^7–14^. The reliable observation of the brain’s dynamic states notwithstanding, mechanistic understanding of the cause and regulation of such dynamics is largely unclear.

Dynamic fluctuations are also ubiquitous in cognitive processing and behavioral performance^15^. Cognition requires the cooperation among distributed networks and is thought to arise as an emergent behavior of the complex brain system^6,16^. Correlational observations have implicated the dynamic brain states in various cognitive processes^17,18^ such as working memory^12,13^ and memory replay^8^. Sustained attention (also known as vigilant attention or tonic alertness) is particularly characterized by substantial fluctuations over time and involves a distributed network of brain areas^19^. Moreover, the neural mechanism underlying sustained attention is thought to fluctuate intrinsically^20^. Specifically, engagement and lapses of sustained attention have been associated with the intrinsic dynamic rivalry of opposing neural networks—the central executive network (CEN; alternatively, the frontoparietal network) and the task-negative network (dominated by the default mode network/DMN)^14,21–23^. In addition, neural synchrony in the alpha frequency has been associated with sustained attention and tonic alertness^20,21,24–28^, and accordingly, alpha-frequency transcranial alternating current stimulation (α-tACS) that augmented alpha power has also been shown to enhance sustained attention^29^. The foregoing thus suggests that sustained attention would provide an ideal model for the study of dynamic brain states, which, conversely, will offer novel systems-level insights into the neural underpinning of this important cognitive process.

Here, we approached these processes by leveraging the inherent spatiotemporal coupling between large-scale network activity and alpha oscillations and manipulating alpha oscillations to perturb brain state dynamics. Particularly, brain state dynamics characterized by DMN activity fluctuations have been repeatedly associated with the presence of strong alpha oscillations in humans^8,30,31^. This potential synergy between the DMN and alpha oscillations aligns with a solid body of conventional (static) studies linking these two processes^32–39^. Recently, experimental manipulations using α-tACS have further established that augmenting alpha oscillations would enhance both DMN fMRI functional connectivity^40^ and DMN alpha oscillations (based on EEG source-level analysis, including both power^31^ and connectivity^40^). That said, to date, such changes have only been measured offline, i.e., following stimulation as aftereffects, which could stem from different processes, precluding a direct inference of the coupling between the DMN and alpha oscillations. Nonetheless, a new rodent study applied simultaneous fMRI and theta-frequency optogenetic modulation, where online, realtime effects provided direct evidence of theta oscillations in driving dynamic brain states^41^, lending credence to such cooperation in humans and compelling the adoption of online brain recordings with brain stimulation in humans to approach such questions.

Therefore, we recorded fMRI simultaneously with high-definition (HD) α-tACS and combined it with a concurrent sustained attention task (the continuous performance task/CPT; **Fig. 1A**). Because sustained attention is modulated by task complexity and cognitive load^19^, we administered the CPT with alternating blocks of high and low load. In addition to the DMN, our analysis incorporated other major cognitive networks (CEN and salience network/SN^13^), as well as the visual network (VN) given the visual task. Using hidden Markov modelling (HMM) of fMRI timeseries from hubs of these large-scale networks^42^, we extracted brain states and characterized their temporal dynamics over the 20-minute task (supplemental **Fig. S1**). After confirming its effects on static brain networks and CPT performance as previously reported^29,31,40^, we tested the hypothesis that α-tACS would modulate the dynamics of brain states (i.e., upregulating task-positive and downregulating task-negative states) in the service of sustained attention.

**Figure 1.**
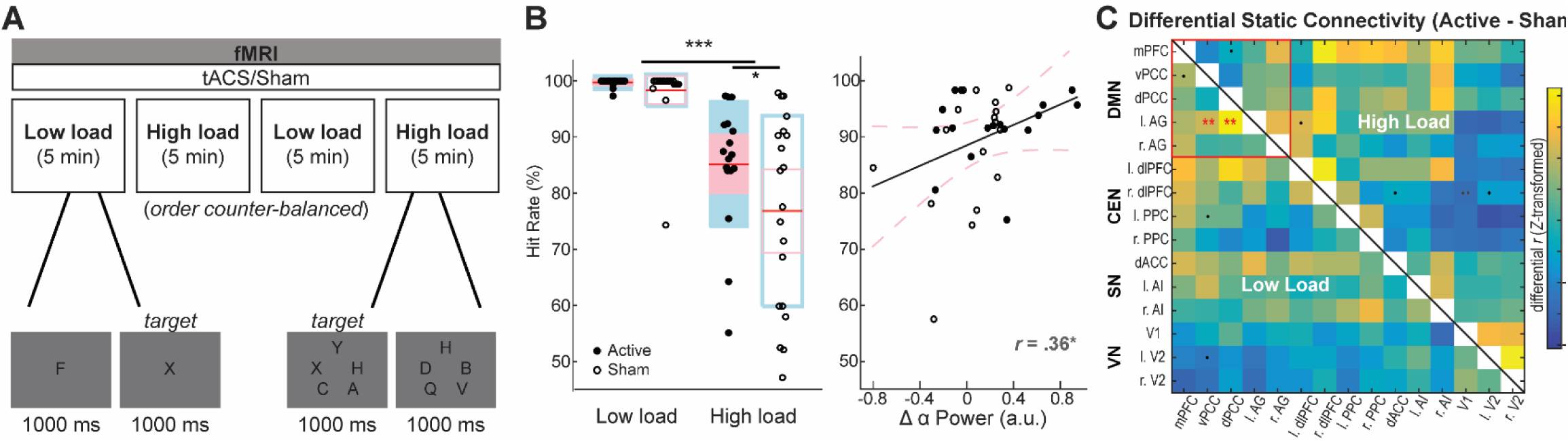
Methods. **A)** Experimental Paradigm. Top: α-tACS or sham stimulation was delivered with simultaneous fMRI recordings while participants performed a sustained attention task. The task (Continuous Performance Task/CPT) consisted of four 5-minute blocks alternating between high and low load conditions (order of the conditions was counterbalanced across participants). Below: Example trials of the task. **B)** CPT performance. Left: The high load condition had lower hit rate than the low load condition in general, but α-tACS (vs. sham control) improved hit rate, particularly in the high load condition. Center red lines represent the mean values, with the pink and blue boxes representing the mean +/- 1.96 SEM and the mean +/- 1.5 SD, respectively. Right: alpha change (Post – Pre) predicted performance (hit rate; collapsed across load levels); a similar correlation was also observed for the high load condition (*Supplemental **Fig. S2***). Active and Sham groups are represented by filled and opened bars and dots, respectively. Dotted pink lines represent 95% confidence interval of least-squares regression line. * = *p* < 0.05; *** *p =* < 0.001. **C)** Conventional network analysis. Differential (Active – Sham) static functional connectivity (Fisher Z-transformed correlations)] matrix for the 15 *a priori* ROIs in low (Lower Left) and high (Upper Right) load conditions. Confirmatory analysis of default mode network (DMN) connectivity demonstrated strengthened connectivity in the DMN for the Active (vs. Sham) group, albeit in the low (but not high) load only. By contrast, significant group effects were absent outside the DMN, highlighting the selective association between the DMN and alpha oscillations. DMN includes mPFC (medial prefrontal cortex), vPCC (ventral posterior cingulate cortex), dPCC (dorsal posterior cingulate cortex), and l/r AG (left/right angular gyrus); CEN (central executive network) includes l/r dlPFC(left/right dorsolateral prefrontal cortex), and l/r PPC (left/right posterior parietal cortex); SN (salience network) includes dACC (dorsal anterior cingulate cortex) and l/r AI (left/right anterior insula); and VN (visual network) incudes V1 and l/r V2. . = *p* < .05, uncorrected; .. = *p* < .01, uncorrected; ** = *p* < .01, FDR corrected.

## RESULTS

### α-tACS target engagement validation

This experiment was a part of a larger study, where participants were randomly assigned to receive 20- minute active or sham HD α-tACS targeting the primary cortical source of alpha oscillations—the occipitoparietal cortex. The α-tACS manipulation was validated in a previous report^40^ based on resting-state EEG before and after the experiment: the Active (vs. Sham) group exhibited significant increase in both posterior alpha power and long-range posterior-to-frontal (P➔F) alpha connectivity (measured with Granger causal/GC connectivity). This change was specific to the alpha frequency and absent in other frequencies. We further conducted a separate, independent experiment with an active control group receiving α-tACS at random frequencies (1-200 Hz), which replicated these effects while ruling out general frequency-non-specific effects. More details are provided in^40^.

### Behavioral effects: **α**-tACS improved CPT performance

During the (active or sham) stimulation, participants completed the 20-minute CPT consisting of two cognitive load levels (low load: a single letter; high load: five letters). Hit rate was submitted to a repeated measures analysis of variance (ANOVA) of Load (high/low) and Group (Active/Sham), which confirmed a load effect: hit rate was significantly higher in the low- than high-load condition, *F*(1,35) = 62.54, *p* <.001, η_p_^2^ = .64. Importantly, as we predicted, there was a Group (i.e., tACS) effect: *F*(1,35) = 3.19, *p* = .042 one-tailed, η_p_^2^ = .08. There was no interaction between Group and Load (*p* = .32). Further linking this behavioral improvement to α-tACS, we confirmed a positive correlation between alpha power increase (Post – Pre tACS) and overall hit rate (*r* = .36, *p* = .039; **Fig. 1B** Right).

We also examined CPT performance based on the variability (the coefficient of variance/CV) of reaction times (RT) throughout the task. A similar ANOVA confirmed the effect of Load, *F*(1,35) = 73.16, *p* < .001, η_p_^2^ = .68; RT was more variable in the high- vs. low-Load condition. Confirming the association between RT variability and sustained attention, higher CV of RT was associated with lower hit rate (*r* = -.57, *p* < .001). However, there were no Group or Load-by-Group effects on RT variability (*p*’s > .66). Lastly, there was no effects of Group on mean RT (*p* = .79), ruling out a speed-accuracy tradeoff elicited by tACS.

### fMRI effects

#### Conventional (static) network analysis

As introduced above, fMRI was recorded during the CPT concurrently with stimulation, and timeseries data was drawn from 15 *a priori* regions of interest (ROIs) encompassing the hubs of DMN, CEN, SN, and VN. For validation, we first confirmed previous findings of α-tACS enhancing resting-state static connectivity in the DMN^40^. Static (conventional time-averaged) DMN connectivity in the low load condition, which, to some extent, approximated a resting state given its minimal cognitive demand, was augmented relative to the pre-tACS baseline in the Active (vs. Sham) group (**Fig. 1C** Lower Left; corrected for multiple comparisons). For comparison, exploratory analyses outside the DMN discovered no connectivity change (that survived correction), highlighting the selective association between the DMN and alpha oscillations. Additionally, in the high load condition (**Fig. 1C** Upper Right), which clearly departed from a resting state, no group effect was significant following correction although at the uncorrected threshold, the right dorsolateral prefrontal cortex (r. dlPFC) exhibited increased connectivity with the anterior cingulate cortex and visual cortex (V1/V2). These exploratory results are not discussed further here; more details are provided in *SI Appendix*.

#### Hidden brain states in sustained attention

For hypothesis testing, fMRI timeseries from the ROIs were submitted to hidden Markov modeling (HMM) to identify dynamic brain states^7,42^. We tested HMMs across a range of one to thirty states. Based on free energy (combined with the “kneedle” method^31,43^), the 8-state HMM was determined as the optimal model (Supplemental **Fig. S1**; see more details in *SI Appendix*). This solution accords with previous studies that converged on HMMs of 8–12 states^7,11,30,42^.

Five of the eight states exhibited moderate-to-strong activation/deactivation (hence denoted as “active” states) in the networks while the other three showed minimal activation/deactivation (i.e., “non-active” states; **Fig. 2**). Based on their specific activation patterns (**Fig. 2**) and combined with the probabilistic time course and transition paths (**Fig. 3**), the five active states were labeled as: 1) “Initiation” state for the clear activation of CEN nodes (and moderate activation of the SN nodes) and the reliable emergence at the onset of (but rarely during) the task blocks (**Fig. 3B**), representative of an alert, active engagement state often observed at the beginning of a task or block; 2) “Task Positive” state for clear activation of the CEN nodes (and moderate activation of the SN nodes) and deactivation of the DMN nodes, consistent with a prototypical task-positive state; 3) “Task Negative” state for the clear activation of the DMN nodes and deactivation of the CEN (barring moderate activation of the left dlPFC) and SN nodes, consistent with a prototypical resting/task-negative state; 4) “Switch” state for the clear activation of dACC (and moderate DMN activation) and deactivation of the CEN nodes, resembling a transition zone between the task-negative and task-positive states (also see transition paths below); and 5) “Distraction” state for the strong right V2 activation, moderate right insula activation, and moderate posterior parietal cortex/PPC activation, which, together, resembled activation of the right-hemisphere dominant ventral attention network^44^. Combined with the DMN deactivation, this state was thus characterized as the Distraction state, potentially induced by visual distractors in the high-load condition (see more discussion below).

**Figure 2.**
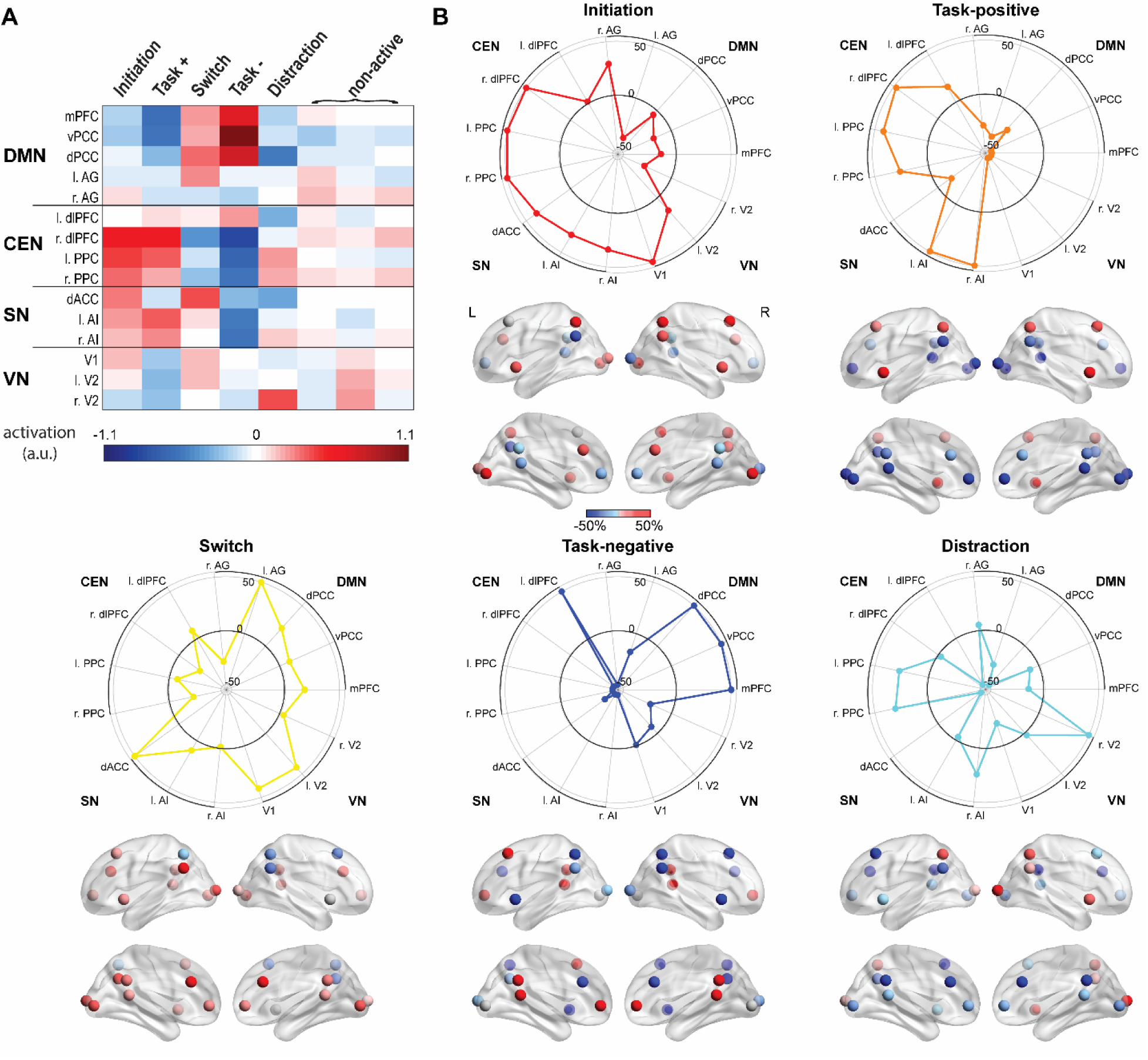
Dynamic brain states during the CPT. **A)** Average activation of each ROI (hub region) of the four networks within each state during the CPT. Positive and negative values indicate higher and lower average BOLD intensities within a given state relative to mean BOLD intensity for the entire CPT, reflecting relative activation and deactivation, respectively. Eight states were identified, including five states with clear ROI activation/deactivation and three states with minimal ROI activation/deactivation. The five (“active”) states were labeled according to their activation/deactivation patterns (as well as transition paths; detailed in Fig. 3). **B)** Normalized activation patterns of the five (“active”) states. Radial plots (Top) and brain models (Bottom) illustrate normalized activation/deactivation levels, i.e., the percent change from baseline of each ROI relative to its maximal activation or deactivation across states. States are color-coded (and henceforth). Top row of brain models is lateral view, and bottom row is medial view. Spheres represent centroids of the anatomical masks of ROIs.

**Figure 3.**
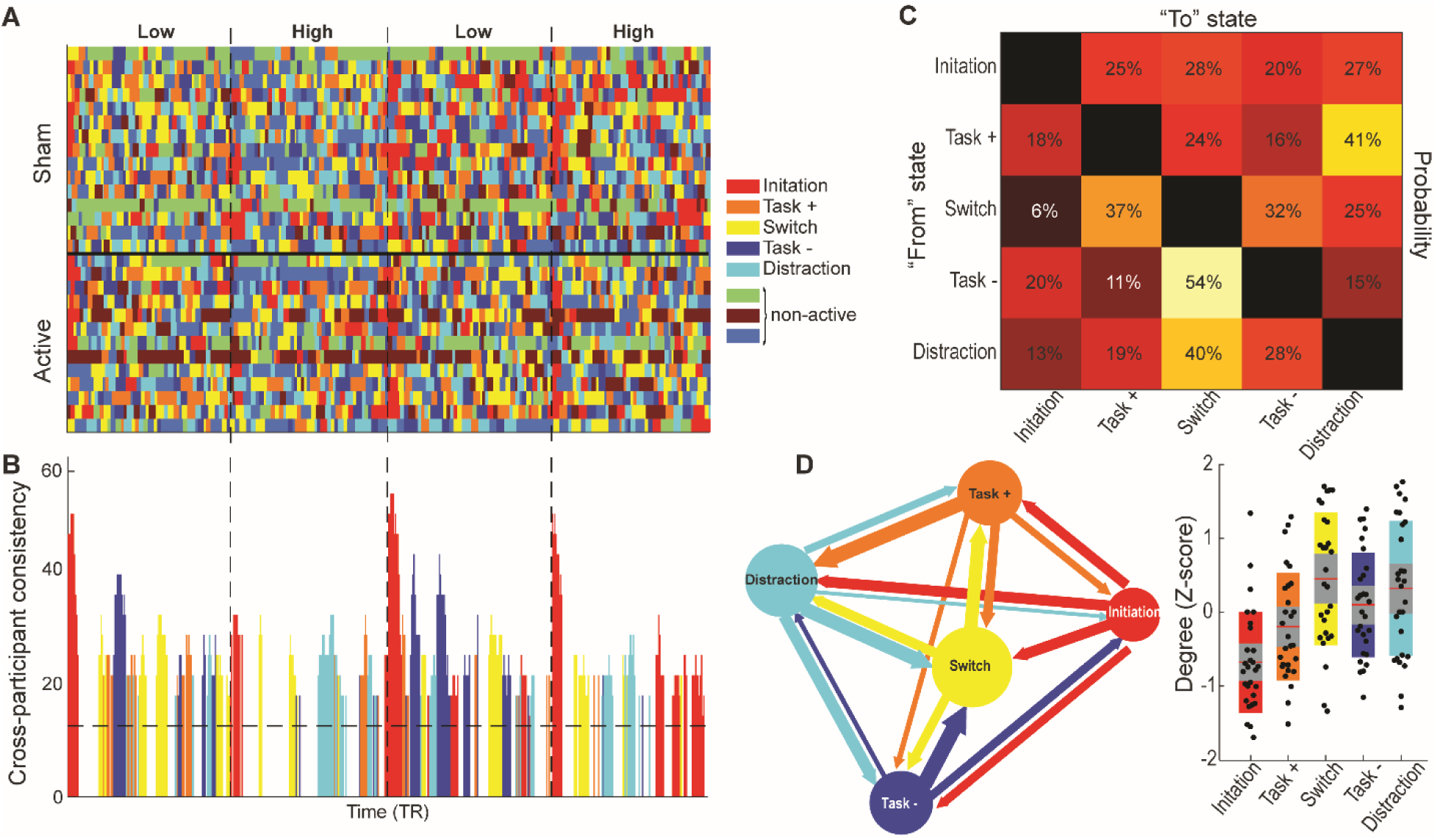
Temporal dynamics of the brain states. **A)** Time courses of all eight states for each participant (based on Viterbi decoding). The onset of each block is marked by a vertical dashed line. **B)** Consistency of state expression across participants for the five (“active”) states. Values indicate the proportion of participants exhibiting the dominant state (based on Viterbi decoding) within a window of 10 TRs^11^. Vertical dashed lines represent onset of task blocks. Horizontal dashed lines represent chance level, i.e., 1 of 8 states (12.5%) being predominantly expressed. **C)** Transition (to and from) probabilities (adjusted for the five active states) for each of the five states, averaged across all participants and task blocks. **D)** (Left) Graph of transition paths between states. Node size represents the fractional occupancy (FO; reflective of overall prevalence of a given state across the duration of the CPT) of each of the five states. Edge thickness represents the transition probabilities. The Switch state was the state with not only the highest FO but also strongest edges. The Task Negative (“Task –”) state tended to transition to the Switch state while the Task Positive (“Task +”) state primarily transitioned to the Distraction state, which then transitioned to the Switch or the Task Negative state. The weakest decile of transition probabilities is not shown. (Right) Centrality (Degree Z-score) of states in the transition graph at individual and group levels. As with edge thickness, the Initiation state had the lowest degree centrality while the Switch state and, to some extent, the Distraction state had the highest degree centrality, reflective of their roles in mediating state transitions. Each dot represents an individual participant, and center red lines represent the mean values, with the grey box and the encompassing box representing the mean +/- 1.96 SEM and the mean +/- 1 SD, respectively.

This activation-based characterization of the states is confirmed by the state transition paths (**Fig. 3C&D**). Particularly, the Switch state appeared to be the transition hub among the active states, serving as the primary transition target for Task Negative state and Distraction state (transition probability = 54%/39%, respectively). While Task Positive state primarily transitioned to Distraction state (reflective of attention deterioration; transitional probability = 41%), its secondary transition target was Switch state (transitional probability = 28%). For outgoing transitions, Switch state transitioned primarily to Task Positive state and Task Negative state (transition probability = 35%/32%, respectively). Finally, the Initiation state exhibited low incoming transition probabilities, akin to its dominance at the beginning of each block. To further qualify and quantify these state transitions, we performed graph theoretical analysis of the transitional probabilities across participants (**Fig. 3D**). The degree centrality index for each state was computed for each subject and submitted to an ANOVA, which showed a significant effect of state (*F*(1,27) = 7.28, *p* < 0.001, η_p_^2^ = .20). Akin to its rather exclusive presence at the beginning of each block, the Initiation state had the lowest degree centrality among all states (FDR *p*’s <0.045). In addition, the Switch state had numerically the highest degree centrality, which was statistically significantly higher than that of the Task Negative state (FDR *p* = 0.025).

#### Temporal dynamics of hidden brain states

We quantified the temporal dynamics of the five active states using two key metrics: fractional occupancy (FO; the percentage of the entire timeseries visited by a state, reflective of the prevalence of that state) and mean lifetime (ML; the average duration of a state visit, reflective of general durability). We then submitted these metrics for each state to separate ANOVAs (Load by Group) to examine the effect of α- tACS on these brain states.

##### Cognitive load modulated state dynamics

The manipulation of cognitive load significantly affected the FO and ML of the Task Negative state. High (vs. low) load reduced both the FO and ML of this state, *F*(1,27) = 15.88, *p* < .001 and *F*(1,27) = 13.24, *p* = .001, respectively (**Fig. 4A&B**). This reduction in both prevalence and durability confirms that the of Task Negative state was generally turned down by high cognitive load. Cognitive load also affected the FO (but not ML) of the Task Positive state. Intriguingly, high (vs. low) load reduced FO of Task Positive state, *F*(1,27) = 4.61, *p* = .041, which, to some extent, may reflect interference caused by distractors in the high load (and indeed, the Distraction state was enhanced in the high load in the Sham group; see below). For other states, the simple effect of Load was not significant (*p*’s > .26).

**Figure 4.**
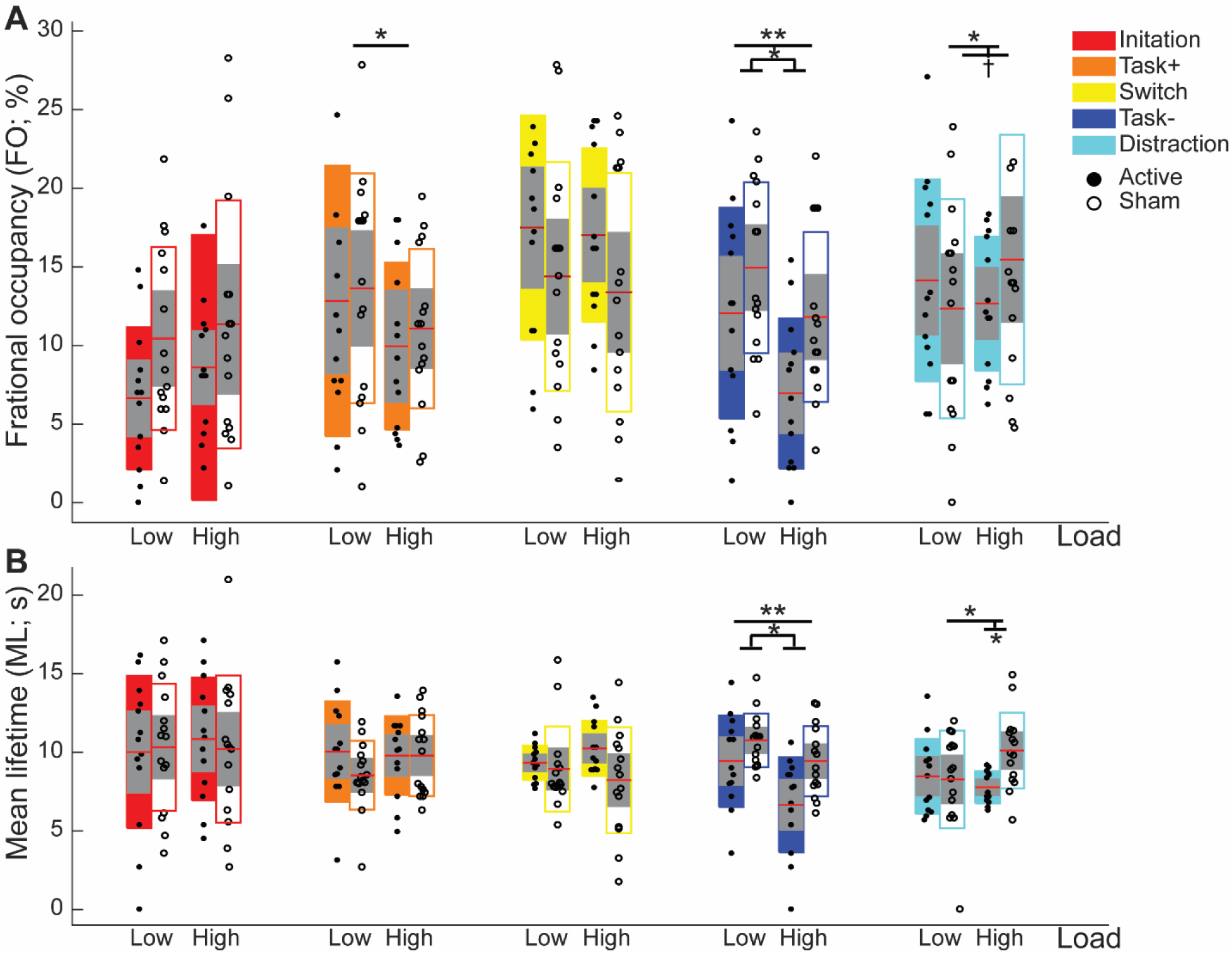
Effects ofα-tACS on temporal dynamics of the states. **A)** Fractional occupancy (FO) and **B)** Mean lifetime (ML) for the five “active” states of the Active (closed circles and boxes) and Sham (open circles and boxes) groups. Cognitive load reduced the FO and ML of the Task Negative (“Task –”) state and the FO of the Task Positive (“Task +”) state. Importantly, α-tACS reduced the FO and ML of the Task Negative state, regardless of load levels. Furthermore, interaction effects of Group and Load on FO and ML of the Distraction state indicate that α-tACS downregulated this state in the high load. Center red lines represent the mean values, with the grey box and the encompassing box representing the mean +/- 1.96 SEM and the mean +/- 1 SD, respectively. * *p* < 0.05; ** *p* < 0.01; † *p* < 0.1.

##### α-tACS modulated state dynamics

The ANOVAs also revealed Group effects on state dynamics. In the Task Negative state, we observed a simple effect of Group on both FO and ML, *F*(1,27) = 4.37, *p* = .047 and *F*(1,27) = 7.36, *p* = .012, respectively, reflecting reduced FO and ML in the Active (vs. Sham) group (**Fig. 4A&B**). Furthermore, in the Distraction state, we observed an interaction between Group and Load on both FO and ML, *F*(1,27) = 4.66, *p* = .040 and *F*(1,27) = 4.98, *p* = .034, respectively. Specifically, as illustrated in **Fig. 4**, high load increased the FO of the Distraction state in the Sham group (*t*(14) = 2.10, *p* = .055), but not in the Active group (*p* = .348). Similarly, ML of the Distraction state was lower for the Active (vs. Sham) group in the high load, *t*(26) = 2.10, *p* = .003, but equivalent for the groups in the low load (*p* = .348). These results suggest that tACS strengthened resistance to distractors in the high load condition. Other states did not exhibit any effects of Group either independently (*p*’s > .09) or interactively with Load (*p*’s > .19). Therefore, α-tACS suppressed Task Negative and Distraction states, especially at high cognitive load.

## DISCUSSION

Applying simultaneous HD-α-tACS and fMRI during a sustained attention task (at low and high load) and employing HMM of fMRI timeseries in major large-scale networks, we uncovered a set of dynamical, functionally important brain states and revealed their responses to cognitive load and alpha modulation. Specifically, we delineated the temporal dynamics of Task Positive state, Task Negative state, and Distraction state, known to facilitate and interfere with sustained attention, respectively, providing mechanistic insights into this important cognitive process and its characteristic fluctuations. Critically, transcranial upregulation of alpha oscillations via α-tACS resulted in the suppression of the interfering states (i.e., Task Negative and Distraction states, especially at high load) and improvement in task performance, highlighting the role of alpha oscillations in regulating dynamics of neural networks and sustained attention. These realtime, online effects thus provide causal insights into the operation of the complex brain system in attention, shedding light on systems-level mechanisms underlying cognition. Finally, that α-tACS selectively modulated two of the dynamic states highlights its dependence on and/or selectivity of ongoing brain states, bearing relevance to future closed-loop applications to optimize tACS.

As detailed in^40^, our α-tACS manipulation led to significant augmentation in alpha oscillations (both alpha power and long-range connectivity). Here, we further demonstrated that it also increased hit rate in the sustained attention task (especially at high load), replicating a prior α-tACS study using a comparable task^29^. Moreover, we found that the degree of alpha augmentation positively predicted hit rates, directly linking α-tACS and the behavioral improvement. These findings add to the growing evidence for the active role of alpha oscillations in cognition (vs. the traditional view of cognitive disengagement or “idling”), particularly for sustained attention and tonic alertness^20,21,24–28^. Our conventional (static) functional connectivity analysis also revealed that α-tACS strengthened static connectivity within the DMN, albeit primarily in the low load condition that approximated a resting state with its minimal cognitive demand. This online effect of α-tACS corroborates previously reported offline (Post – Pre) resting-state connectivity increase in the DMN^40^, ruling out confounds such as rebound effects for the offline finding and highlighting a direct, selective, and enduring effect of α-tACS on intrinsic DMN connectivity.

Our HMM analysis further identified five functionally relevant brain states during the sustained attention task and provided a cohesive depiction of their temporal dynamics. Critically, α-tACS modulated these dynamics, particularly in the high load condition. In fact, as illustrated in **Fig. 1B** Left, behavioral improvement was primarily present at this load (*t*(33.22) = 1.79, *p* = .042 one-tailed), which correlated with alpha power increase (*r* = .36, *p* = .037), but failed to reach significance in the low load condition (*t*(19.75) = 1.07, *p* = .149 one-tailed; see details in *SI Appendix* and *Supplemental **Fig. S2***). Specifically, similar to an earlier study with interleaved low- and high-load blocks^13^, we uncovered a state that emerged at the onset of every block, which was characterized by strong CEN and moderate SN activation, akin to the initiation (and transition) of the task (and load). In addition, attention engagement and lapses have been associated with the involvement of the CEN (aka, frontoparietal network) and the DMN, respectively^22^, and indeed, we not only identified a Task Positive state (characterized by CEN activation and DMN deactivation) and a Task Negative state (characterized by DMN activation and CEN deactivation) but also revealed their responsiveness to cognitive load. Particularly notable, the load effect on the DMN-dominant (Task Negative) state (i.e., higher FO/ML at low than high load) corroborated the notion that the DMN is activated during rest and low-load tasks and de-activated during effortful tasks^45^.

Furthermore, the brain vacillated between these two states via a Switch state. In keeping with this, the Switch state was more visited than other states, i.e., with greater FO than all other states (*p*’s < .02) except for the Distraction state (*p* = .13; **Fig. 3C&D**). This frequent transition between Task-Positive and Negative states accords with constant fluctuations characteristic of sustained attention. It has been postulated that the vacillation between Task Positive and Task Negative states may reflect an adaptive process to prevent over-engagement of the CEN and over-disengagement of the DMN, which could undermine performance^22^. Therefore, attention fluctuation (especially, over an extended period of sustained attention) may reflect the rhythmicity of attention that is cognitively beneficial^46^. Conversely, the absence of such fluctuating (or even labile) states (and therefore the perpetuation of a certain state) would cause avalanches of the complex brain system, resulting in cognitive impairments and even underpin neuropsychiatric disorders^47^. For instance, entanglement between the DMN and SN^14,48^ and low neural variability in all four networks^49^ in attentional disorders (e.g., attention deficit hyperactivity disorder) could reflect disruptions in the dynamical fluctuation of brain states, underpinning its attentional impairments. In keeping with that, we observed that while improving the CPT accuracy, α-tACS did not reduce variability (CV of RT) of the performance. Consistently, we observed no effect of α-tACS on the transition rate of the brain states (*p*’s >.10). That is, while adjusting the balance between functionally beneficial and detrimental states (more discussion below), α-tACS preserved neural fluctuation (or rhythmicity) throughout the task.

Finally, we observed a Distraction state that was particularly pronounced at high load (characterized by the presence of distractors) in the Sham group. Interestingly, the Task Positive state was prone to transition into this Distraction state, which further transitioned into the Switch state or defaulted into the Task Negative state (**Fig. 3C&D**; **Fig. 4**). This suggests that attention lapses could arise as the Task Positive state is hijacked by distracting input. Importantly, both the Task-Negative and Distraction states were suppressed by α-tACS (**Fig. 4**), in keeping with the behavioral improvement it induced. It is also worth noting that efficacy of α-tACS has been shown to be state-dependent^50^, and a study examining its aftereffect (during post-tACS resting-state recordings) on dynamic brain states indicated that it primarily affected a DMN-dominant state^31^. Therefore, the current finding underscores this state-dependent quality of alpha stimulation and its close association with DMN functioning, and thus promotes the application of closed-loop α-tACS to fully capitalize upon its neuromodulatory capacity.

The static and dynamic effects of α-tACS together causally illuminate how the complex brain system operates on its primary architectures—large-scale networks and neural synchrony. Specifically, we surmise that at rest, alpha oscillations upkeep intrinsic DMN integrity whereas during task, they regulate the dynamic balance (“on” and “off”) of the major large-scale networks that are conducive or disruptive to task performance. This notion resonates with increasing recognition of the multifaceted, sometimes paradoxical, functions of alpha oscillations^27,51^. The former (supporting intrinsic DMN functioning at rest) features alpha oscillations as a “long-range communicator”^27,51^. Specifically, as reported previously^40^, the strengthening of intrinsic DMN connectivity via α-tACS was mediated by frontal-posterior alpha synchrony (indexed by posterior-to-frontal alpha-frequency Granger causality) but not by local alpha activity (indexed by alpha power). In comparison, the latter (regulating brain state dynamics during task, particularly sustained attention) exemplifies local modulation by alpha oscillations as a “sensory inhibitor” (that suppresses distracting information and thus a Distraction state) and a “vigilance maintainer” (that fends off a Task Negative state)^27,51^. In keeping with this latter function, we observed that not only task performance but also the durability (i.e., ML) of Distraction state was predicted by posterior alpha power (*r* = -.42, *p* = .037; see *SI Appendix*). Moreover, the strong right V2 activation in the Distraction state aligns with the right-hemisphere lateralization of local alpha inhibition of the sensory cortex^52^. Consistent with previous combined EEG-fMRI studies^53,54^, this transient activation of the right V2 in the Distraction state likely reflects evasion from alpha inhibition (i.e., failed sensory gating or filtering), resulting in increased response to distracting visual stimuli. Together, behavioral and neural (static and dynamic) effects α-tACS coalesce to highlight and harmonize the complex functions of alpha oscillations.

In summary, current findings provide new insights into the dynamical organization of the complex brain system that underpins cognition. Specifically, it presents a complex system perspective of the mechanism underlying sustained attention: the engagement of a tug-of-war between Task Positive and Task Negative brain states (along with resulting frequent switching between them) and the interception of the Task Positive state by distractors. Critically, alpha oscillations play a modulatory role in such dynamics of brain states, effectively shifting the balance in the tug-of-war (favoring beneficial over detrimental states). Consequent to the fine tuning of brain state dynamics, sustained attention performance improves.

## Materials and Methods

### Participants

Forty-one healthy volunteers (24 females, 20.8 ± 3.2 years of age) participated in the experiment as a part of a larger study. Participants reported no history of neurological or psychiatric disorders, current use of psychotropic medications, and had normal or corrected-to-normal vision. Participants were randomly and blindly assigned to the Active group (*n* = 21) or the Sham group (*n* = 20). Two participants (Active *n* =2) terminated the experiment due to discomfort in the MRI scanner. fMRI data were collected during the CPT from twenty-nine participants, and one Sham participant was excluded from analysis due to excessive motion, resulting in 28 participants for fMRI analysis (Active *n* = 13, Sham *n* = 15). Two Active participants were excluded for behavioral recording errors and failure to follow instructions, respectively, resulting in 37 participants for behavioral analyses (Active *n* = 17, Sham *n* = 20). The two groups did not differ in age or gender (*p*’s > .5). Experimental protocol was approved by Florida State University’s Institutional Review Board.

### Experimental Design

Participants performed a sustained attention task for 20 mins while fMRI data was collected and tACS or sham stimulation was delivered (**Fig. 1A**). The task was administered before and after resting-state EEG and fMRI data acquisition (reported in^40^).

#### Continuous Performance Task (CPT)

The continuous performance task (CPT) has been widely used to study sustained attention^55,56^. In this study, the CPT task included two conditions, a low-load condition (a single letter at center of the screen) and a high-load condition (5 equidistant letters encircling a central fixation point, and in the target trial, the target letter was presented with 4 other letters, i.e., distractors; **Fig. 1A**), each presented in two blocks in alternating orders counterbalanced across participants and groups. Each block consisted of 300 trials, each lasting 1000 ms for a total of five minutes. Participants were instructed to respond as quickly as possible via button press when the letter “X” appeared on screen. Target trials occurred on 12.5% of the trials in each block.

#### tACS

Alpha-frequency stimulation was administered for the entire 20 minutes of the CPT. A ±2 mA sinusoidal current oscillating at 10 Hz was applied using an MR-compatible High-Definition (HD) tACS system in a 4 × 1 montage over midline occipitoparietal sites, which were selected to maximally target the primary cortical source of alpha oscillations—occipitoparietal cortex. Sham stimulation was similarly applied, but the current was on for the first and last 30 seconds of the 20-minute task. Blindness of group assignment were confirmed through a systematical assessment. More details are provided in^40^.

### MRI Acquisition and Preprocessing

Gradient-echo T2-weighted echoplanar images were acquired on a 3T Siemens Prisma MRI scanner using a 64-channel head coil with axial acquisition. Imaging parameters were TR/TE: 1800/22.40 ms; flip angle: 40°, field of view 212 mm, slice thickness 1.8 mm, gap .45 mm; in-plane resolution/voxel size 1.8 × 1.8 mm; multiband acceleration factor = 2; GRAPPA acceleration factor = 2. A high-resolution (.9 × .9 × .9 mm3) 3D-MPRAGE T1 scan was also acquired. Imaging data were preprocessed using SPM12, including slice-time correction, spatial realignment, and normalization using Diffeomorphic Anatomical Registration Through Exponentiated Lie algebra (DARTEL). We then submitted the timeseries to DPARSFA toolbox^57^ for additional preprocessing, including (1) mean centering and whitening of timeseries; (2) temporal bandpass (.01-.08 Hz) filtering; and (3) nuisance regression with 24 variables (six head motion parameters each from the current and previous scan and their squared values). Details of imaging parameters and preprocessing protocols were described in^40^.

#### Regions of Interest (ROIs)

The three major cognitive neural networks—DMN, CEN, and SN—and the visual network (VN) were included. A total of 15 regions of interest (ROIs) representing hub regions of the four networks were included. Specifically, the DMN ROIs included midline (medioprefrontal cortex/mPFC, ventral and dorsal posterior cingulate cortex/vPCC & dPCC) and lateral (left and right angular gyrus/AG) hubs of the DMN; CEN ROIs included the left/right dorsolateral prefrontal cortex (dlPFC) and the left/right posterior parietal cortex (PPC); the SN ROIs included dorsal anterior cingulate cortex (dACC) and left/right anterior insula (AI); and the VN ROIs included V1 and left/right V2. The CEN SN, and frontal and lateral DMN ROIs were defined by the Willard Atlas^58^. The PCC subdivisions were individually defined using the Brainnetome Atlas^59^. The VN ROIs were defined by a probabilistic atlas of the visual cortex^60,61^.

#### Hidden Markov Modeling (HMM)

Preprocessed fMRI timeseries from the entire task were drawn from the ROIs and submitted to modeling via the HMM-MAR toolbox (https://github.com/OHBA-analysis/HMM-MAR). From our models generated with between 1 and 30 states, we determined that the 8-state model optimally represented the data. The model fit was indexed by the free energy, Akaike Information Criterion (AIC), Bayesian information criterion (BIC), and integrated complete likelihood (ICL) metrics. More details are provided in *SI Appendix* and *Supplemental **Fig. S1***.

#### Graph theoretical analysis

Graph theoretical analysis was performed on transition probabilities of the five “active” states using the Brain Connectivity Toolbox (https://github.com/brainlife/BCT) and visualized in Gephi^62^. To examine centrality of the five states, we calculated degree Z-score for each of the states in each participant and submitted the values to statistical analysis.

### Statistical analysis

Target engagement of tACS (increased alpha power and P➔F alpha connectivity) was validated previously^40^. We then confirmed the previously reported effect of tACS in improving sustained attention^29^ using conducting repeated measures analyses of variance (ANOVAs) of Load (high/low) and Group (active/sham) on CPT accuracy and coefficient of variance (CV) of RT, respectively. For this confirmatory analysis, statistical significance was set at *p* < .05 one-tailed. Other than confirmatory analyses on the DMN connectivity (see *SI Appendix)*, we applied false discovery rate/FDR correction on tests for all other connections. For hypothesis testing (regarding the dynamics of brain states), we conducted similar ANOVAs (Load by Group) on the FO and ML on the identified states, respectively. Statistical significance was set at *p* < .05 two-tailed.

## Acknowledgments

This research was supported by the National Institutes of Health grants (R01MH132209 and R01NS129059 to W.L.) and the FSU Chemical Senses Training (CTP) Grant Award T32DC000044 (K.C.) from the National Institutes of Health (NIH/NIDCD).

