## Supplemental information for "Transcranial stimulation of alpha oscillations modulates brain state dynamics in sustained attention"

**This PDF file includes:**

Supplementary text

Figures S1, S2, and S3

SI References

**Supplementary Methods**

**Conventional (static) network analysis**

Preprocessed fMRI timeseries for the high- and low-load blocks were drawn from each of the 15 ROIs (including the hub nodes of DMN, CEN, and SN, in addition to V1 and bilateral V2 that constitute the visual network/VN) and submitted to Pearson’s correlation analysis to construct a 15x15 correlation matrix for each condition for each participant. The pair-wise correlation coefficients were then Fisher Z-transformed and corrected for baseline connectivity (by subtracting Pre-tACS resting-state connectivity) before submission to statistical analyses.

**Hidden Markov Modeling (HMM)**

The HMM assumes that dynamic fluctuations in multivariate timeseries across a set of brain regions (or networks) can be characterized by a sequence of discrete hidden brain states, which transition and recur over time at a certain time-invariant probability. This approach has been validated in both rest and task^2^. We inferred the HMM using the publicly available HMM-MAR (Hidden Markov Model-Multivariate Autoregressive) toolbox (<https://github.com/OHBA-analysis/HMM-MAR>^3^), which represents each state by a multivariate Gaussian distribution and provides estimates of the parameters of the state distributions and the probabilities of each state to be active at each time point. Following previously established methods, we submitted the 15 (ROIs) x 556 TRs data matrix across 28 subjects to Gaussian HMM inference using 500 training cycles and 50 repetitions. Both the mean (within each ROI) and the covariance (between ROIs) of the distribution were considered during training. State time sequences were obtained using the Viterbi decoding algorithm.

We tested HMMs across a range of 1 to 30 states and determined the best number of states based on free energy combined with the “kneedle” method^4,5^. As illustrated in Supplemental **Fig. S1**, the benefit of additional states sharply decreased (akin to a knee) and became minimal beyond 8 states, coinciding with previous studies that converged on HMMs of 8–12 states^2,6-8^. We also examined the Akaike information criterion (AIC) metric, the Bayesian information criterion (BIC), and the integrated complete likelihood (ICL)^9^ ,which converged with the free energy metric. The 8-state HMM was thus chosen as the optimal model. Based on this HMM, we then derived (i) fractional occupancy (FO) and mean lifetime (ML) for each state and (ii) state transition matrices for the probability/path of state transition.

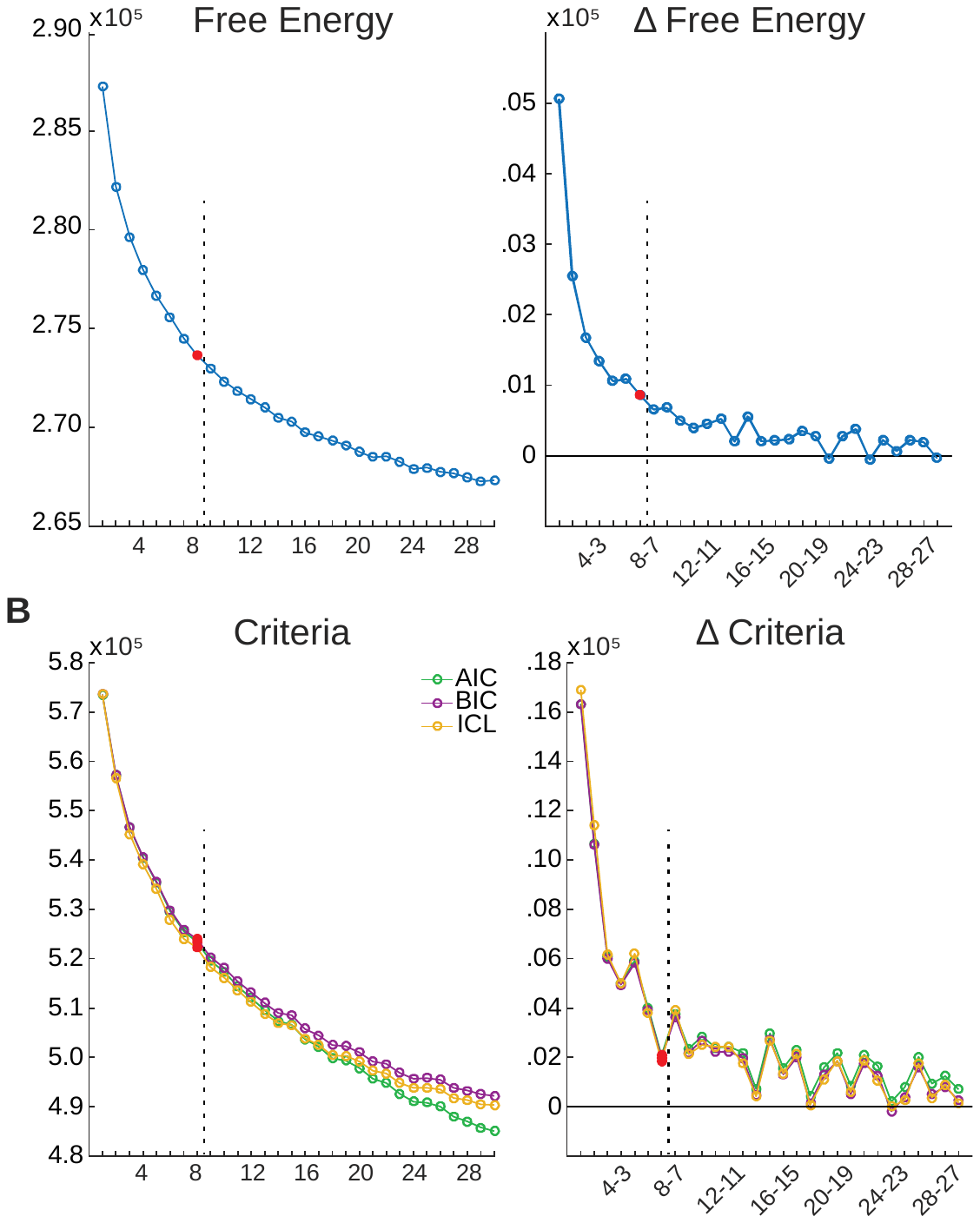

**Supplemental** **Fig. S1. HMM models.** HMMs were generated between 1 and 30 states. A) Free energy (Left) and change in free energy (Right) as a function of states indicate a largely monotonical decrease as the number of states increased. A knee emerged around 8 states (red circle), which was further followed by plateaued reduction in free energy. B) The Akaike information criterion (AIC) metric, the Bayesian information criterion (BIC), and the integrated complete likelihood (ICL) indicated similar decay patterns to the free energy.

**Supplementary Results**

**Behavioral effects: α-tACS primarily improved high-load CPT performance**

As illustrated in **Fig. 1** (main text), effects of α-tACS on hit rate were primarily present in the high load condition, which was confirmed by simple contrasts in each cognitive load. Hit rate was improved in the active group (relative to sham) for the high load (*t*(33.22) = 1.79, *p* = .042 one tailed) but not the low load (*t*(19.75) = 1.07, *p* = .149 one tailed), presumably due to a ceiling effect in the low load. Moreover, as illustrated in **Fig. S2**, there was a positive correlation between resting alpha power change (Post – Pre tACS) and hit rate in the high load (*r* = .36, *p* = .037 two tailed), but no relation in the low load (*p* = .244). These results suggest that α-tACS improves sustained attention primarily when greater cognitive engagement is required. No relationships for RT mean or variability were identified with tACS group or alpha power change (*p*’s > .392), regardless of cognitive load, ruling out the possibility that the α-tACS effects were merely due to a speed-accuracy tradeoff.

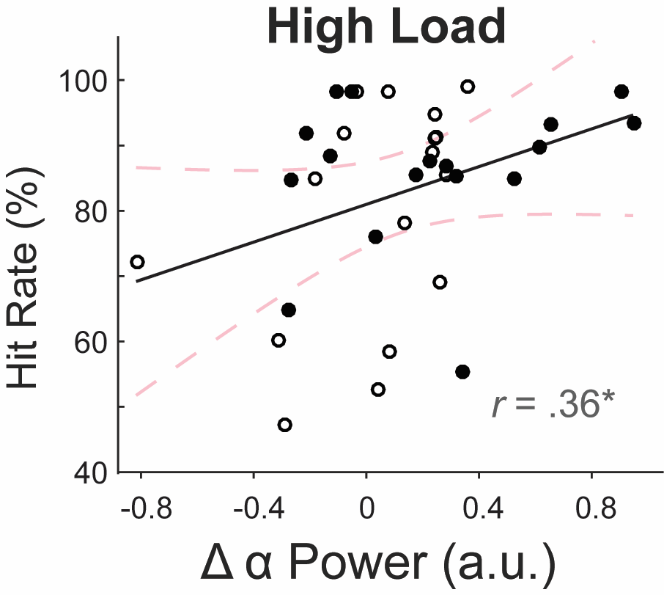

**Supplemental** **Fig. S2.** Resting alpha power change (Post – Pre) positively correlated with the hit rate in the high load. Active and Sham groups are represented by filled and opened dots, respectively. * = *p* < .05.

**Neural effects: Correlation of alpha oscillations with dynamics of brain states**

We explored the association of alpha power change and the dynamics metrics for the two states (the Task Negative state and Distraction state) that were modulated by α-tACS. As illustrated in supplemental **Fig. S3**, the ML of Distraction state in the high load was negatively predicted by alpha power change, linking alpha oscillations with the suppression of this state (*r* = -.42, *p* = .037).

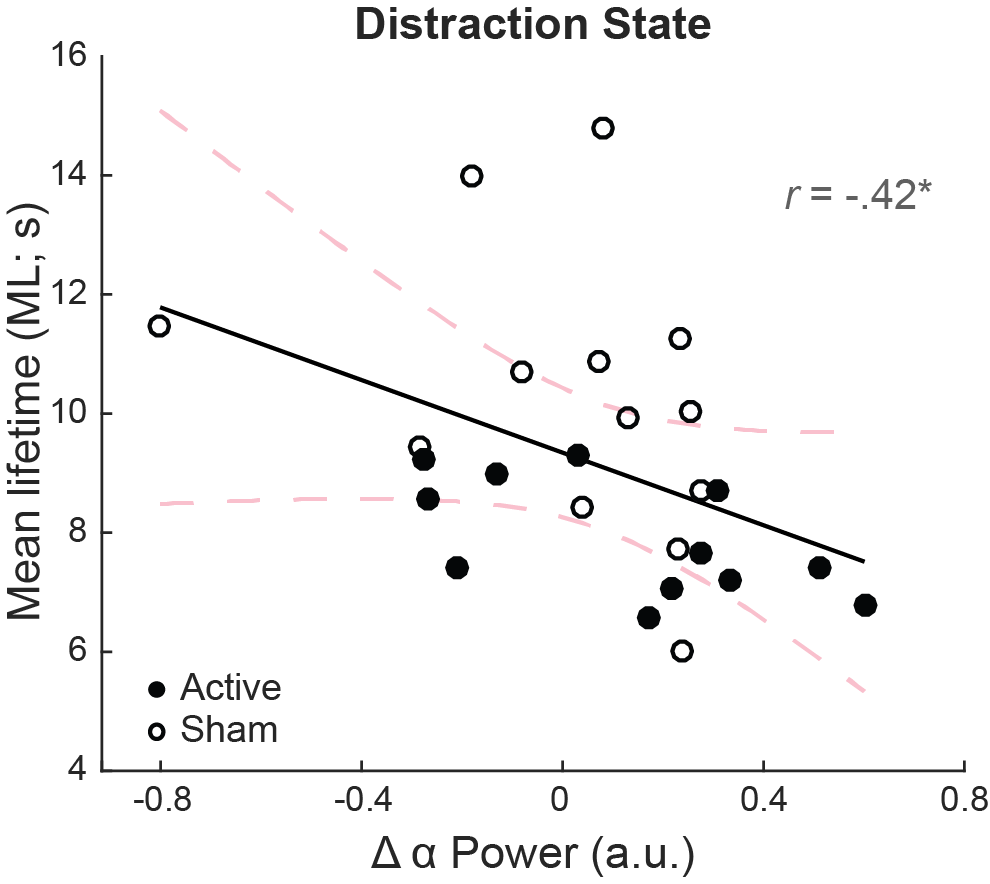

**Supplemental** **Fig. S3.** Alpha power change (Post – Pre) negatively correlated with the ML of high-load Distraction state. Active and Sham groups are represented by filled and opened dots, respectively. * = *p* < .05.
